# The Metacognitive Sensitivity of Verbal Expressions of Confidence in a Perceptual Decision

**DOI:** 10.64898/2026.05.13.724887

**Authors:** Facundo Alvarez Heduan, Ariel Zylberberg

## Abstract

We study how confidence in perceptual decisions depends on whether it is communicated verbally (e.g., “very likely”) or numerically (e.g., “80% certainty”). We find that verbal expressions more reliably distinguish correct from incorrect choices than numerical reports, challenging the common assumption that numerical probabilities provide more precise representations of uncertainty. Additionally, in a dyadic decision-making task in which participants can revise their initial reports based on a partner’s choice and expressed confidence, verbal and numerical reports are equally effective in supporting accurate revisions of initial judgments. Together, these results underscore the effectiveness of verbal expressions as a means of conveying decision confidence.

## Introduction

Decision confidence reflects a decision-maker’s belief that their choice is correct. Computational and normative theories often formalize this belief within a probabilistic framework, interpreting confidence as the posterior probability that a decision is correct given the available evidence (Joyce, 1998; Rosenkrantz, 1981; Pouget et al., 2016; Hangya et al., 2016; Zylberberg et al., 2018). This interpretation naturally motivates numerical confidence reporting, since it allows subjective beliefs to be elicited on scales that can be treated as probabilities or monotonic transformations thereof (e.g., Lichtenstein and Fischhoff, 1982; Lawson et al., 2023).

Despite this normative appeal, it remains unclear whether numerical formats are the most effective way to capture and communicate subjective confidence. Verbal expressions of uncertainty (e.g., “likely,” “maybe”) provide a natural alternative: they are ubiquitous across languages, predate formal probability theory, are acquired early in development, and are widely used in everyday communication (Hacking, 2006; Diuk et al., 2012; Zimmer, 1983; Wallsten et al., 1993).

Most studies comparing verbal and numerical expressions of uncertainty has focused on judgments about external events. Representative examples include predictions of geopolitical outcomes (Beyth-Marom, 1982) and sports forecasts (Erev and Cohen, 1990). In such cases, uncertainty is typically well characterized within a frequentist framework: outcomes are independent of the forecaster’s internal state and accuracy can often be improved through aggregation across individuals. Within this literature, numerical expressions are generally found to yield more accurate and consistent judgments than verbal ones. This advantage has been attributed to substantial inter- and intra-individual variability in the interpretation of verbal expressions, as well as to the difficulty of performing arithmetic operations over linguistic representations (Budescu et al., 2014; Brun and Teigen, 1988; Lichtenstein and Newman, 1967; Ott, 2021; Mauboussin and Mauboussin, 2018; Mosteller and Youtz, 1990; Mandel et al., 2021; Mislavsky and Gaertig, 2022).

By contrast, confidence in one’s own decisions is inherently self-referential and depends on internal evidence that may not be naturally represented as precise numerical quantities. Instead, such confidence may be encoded in more qualitative or graded linguistic forms, potentially making verbal expressions comparatively well suited for this domain. However, empirical work directly testing this possibility is limited, and has often relied on complex paradigms such as eyewitness identification, where verbal and numerical reports show comparable predictive validity (Mansour, 2020; Smalarz et al., 2021). This leaves open whether the relative advantage of numerical over verbal formats extends to decision confidence.

Here, we compare verbal and numerical confidence reports in a simple perceptual decision-making task. Perceptual tasks allow for precise control over difficulty and are supported by well-established computational models of confidence (Gold and Shadlen, 2007; Pouget et al., 2016; Fleming, 2024; Fleming and Daw, 2017). We evaluate how well each reporting format discriminates between correct and incorrect decisions and test their impact in a dyadic setting, where participants can revise their choices based on a partner’s expressed confidence (Yaniv, 2004; Koriat, 2012).

## Results

### Task Overview and Experimental Conditions

Participants (N = 8,308) had to decide whether a circular aperture contained more red or blue dots (Fig. 1A). Decision difficulty was controlled by the “color strength”—the proportion of dots in the majority color. On each trial, the color strength was set to one of the following values: 52.5%, 55%, 60%, or 70%. Participants also reported their confidence in the accuracy of their decision (Fig. 1B). Some participants used a numerical scale ranging from 50 to 100, where they indicated the probability that their choice was correct by adjusting a slider (Fig. 1B). Others selected from a list of verbal expressions the one that best described their confidence (Fig. 1C-D).

**Figure 1.**
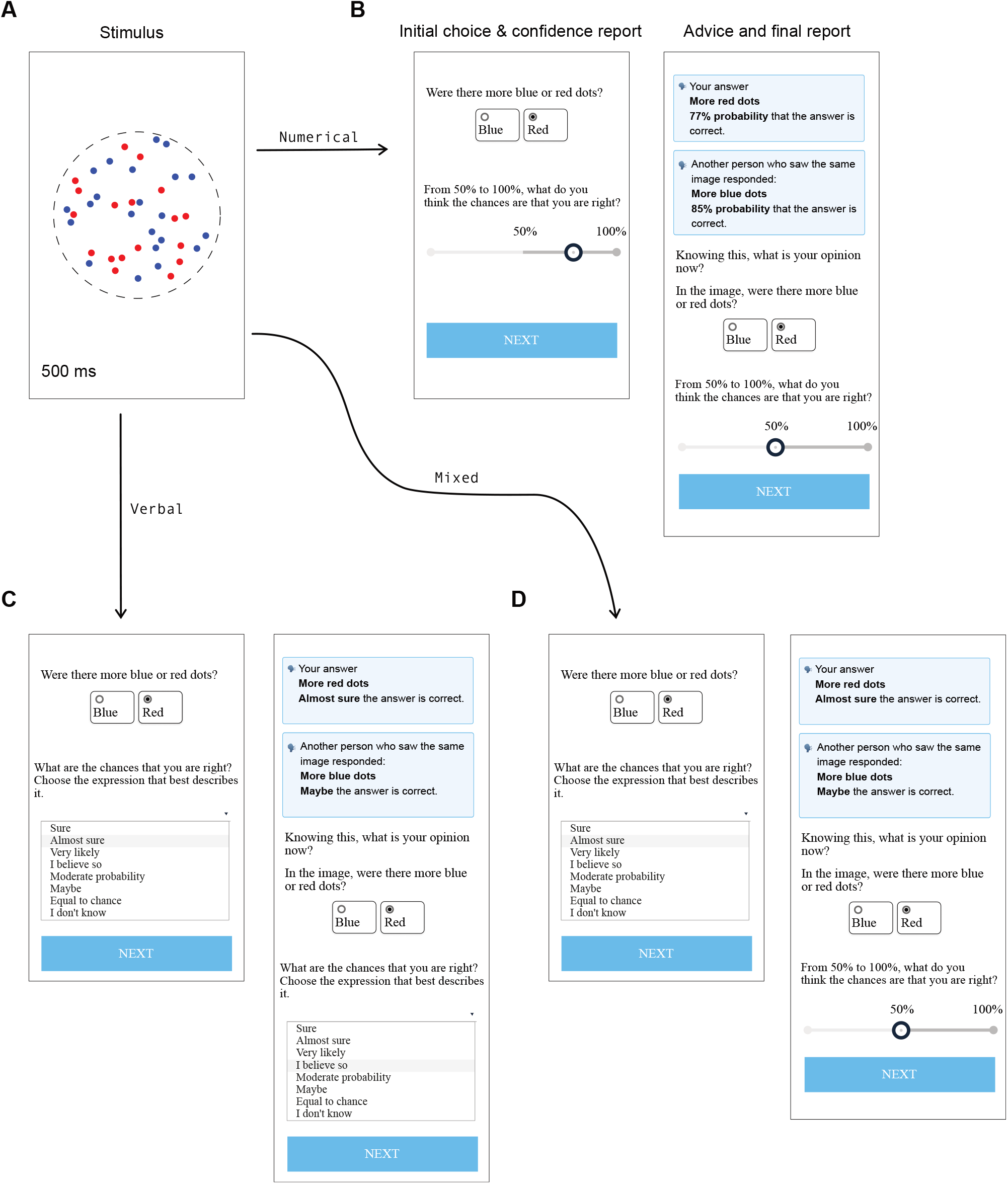
Color Discrimination Task. Sequence of events in a trial. The figure panels closely resemble what participants observed when performing the task. **(A)** Example of the color stimulus. Participants decided whether the stimulus contained more blue or red dots. Decision difficulty was manipulated through *color strength*, defined as the proportion of dots drawn in the majority color. The stimulus was displayed for 500 ms, after which participants (i) reported their choice (blue/red) and their confidence, (ii) observed the choice and confidence reported by another randomly selected participant who had seen a stimulus with the same color strength and majority color, and (iii) provided their final choice and confidence report. **(B)** Numerical Only condition: All confidence reports—initial, advice, and final—were numerical. Participants used a slider on a scale ranging from 50% to 100% to indicate their confidence, where 50% indicates chance-level accuracy (i.e., guessing). **(C)** Verbal Only condition: All confidence reports were verbal, selected from a predefined containing the eight verbal expressions indicated in the figure panel. **(D)** Mixed condition: Initial and advice confidence reports were verbal, while the final confidence report was numerical.

After making their initial choice and confidence report, participants were presented with “advice”—the choice and confidence of another randomly selected participant who had seen a stimulus with the same majority color and strength. Participants then provided their final choice and confidence, potentially revising their initial responses based on the advice. Each participant completed two trials per color strength (one with more blue dots and one with more red dots), totaling eight trials per participant.

The key manipulation of the experiment is the format used to report confidence. Participants were assigned to one of three conditions (Numerical Only, Verbal Only and Mixed), which varied in whether confidence was reported verbally or numerically:

|  |  |
| --- | --- |
| Numerical Only | Initial confidence, advice confidence, and final confidence were reported numerically (Fig. 1B). |
| Verbal Only | Initial confidence, advice confidence, and final confidence were reported by selecting a verbal expression from a list (Fig. 1C). |
| Mixed | Initial confidence and advice confidence were reported verbally, and final confidence was reported numerically (Fig. 1D). |

The Numerical Only, Verbal Only, and Mixed conditions were completed by 2,740, 2,856, and 2,712 participants, respectively.

Before the main experiment, participants completed a mapping task (Fig. 2). They were presented with 16 verbal expressions of uncertainty and were asked to assign to each one the numerical value (0–100% probability scale) that best captured its meaning (Fig. 2A). As shown in Fig. 2B, the numerical values associated with each expression displayed considerable variability across participants, consistent with findings from many previous studies (for a review, see Dhami and Mandel, 2022).

**Figure 2.**
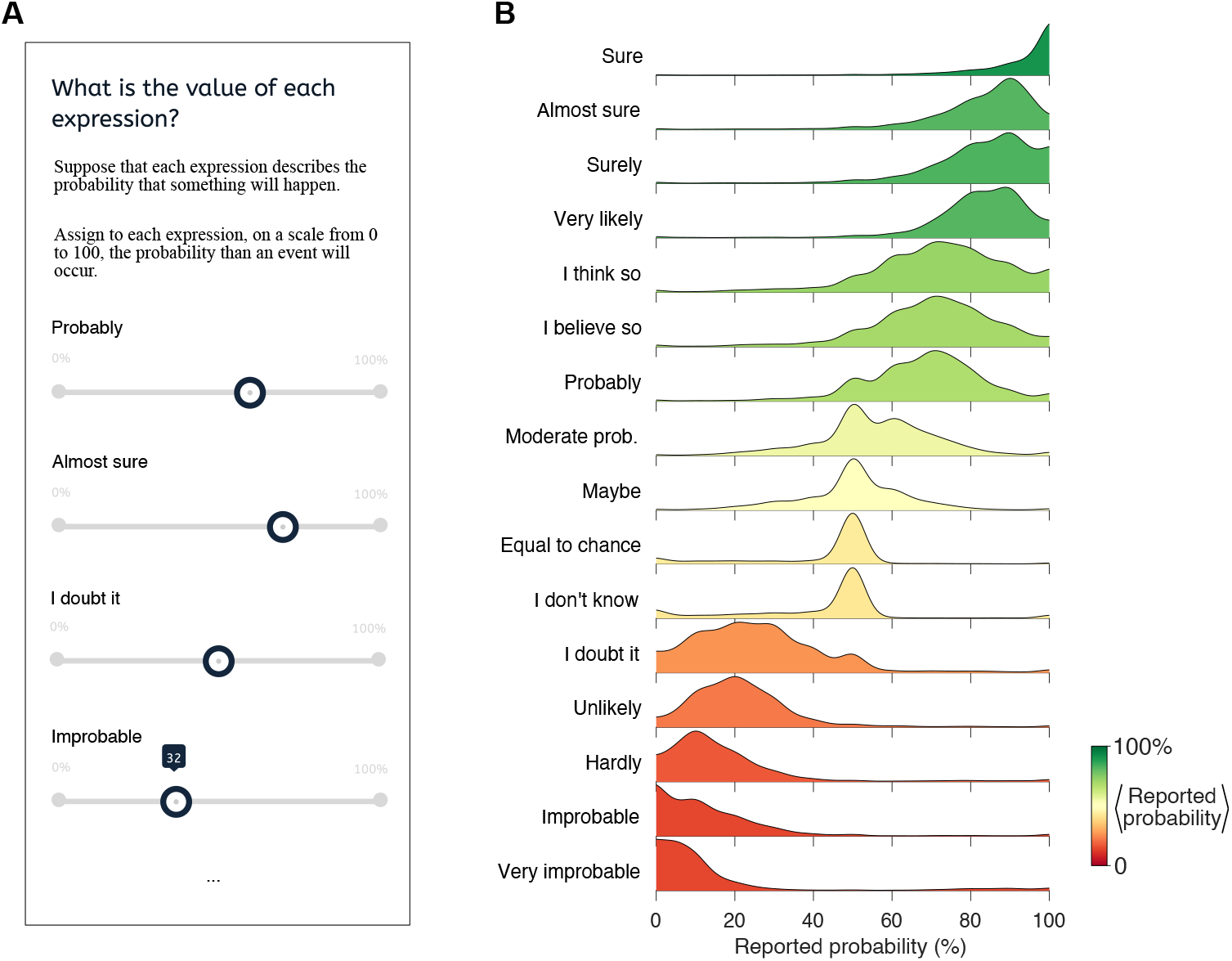
Mapping Task. **(A)** Participants assigned numerical probabilities (0–100 range) to 16 verbal expressions of certainty presented in random order. **(B)** Distribution of numerical values associated with each expression, sorted by mean value. Gaussian kernel smoothing was used to estimate probability densities. Colors represent the average value assigned to each expression.

### Relation Between Color Strength, Accuracy, and Confidence

Initial reports revealed a consistent relation between color strength, decision accuracy, and confidence. Higher color strength led to higher accuracy (Fig. 3A) (*p* < 10^−8^; ℋ_0_: *β*_1_ = 0; Eq. 3, likelihood-ratio test). Choice accuracy was unaffected by whether confidence was reported verbally or numerically (Fig. 3A) (*p* = 0.62; ℋ_0_ : *β*_2,3_ = 0; Eq. 3, likelihood-ratio test).

**Figure 3.**
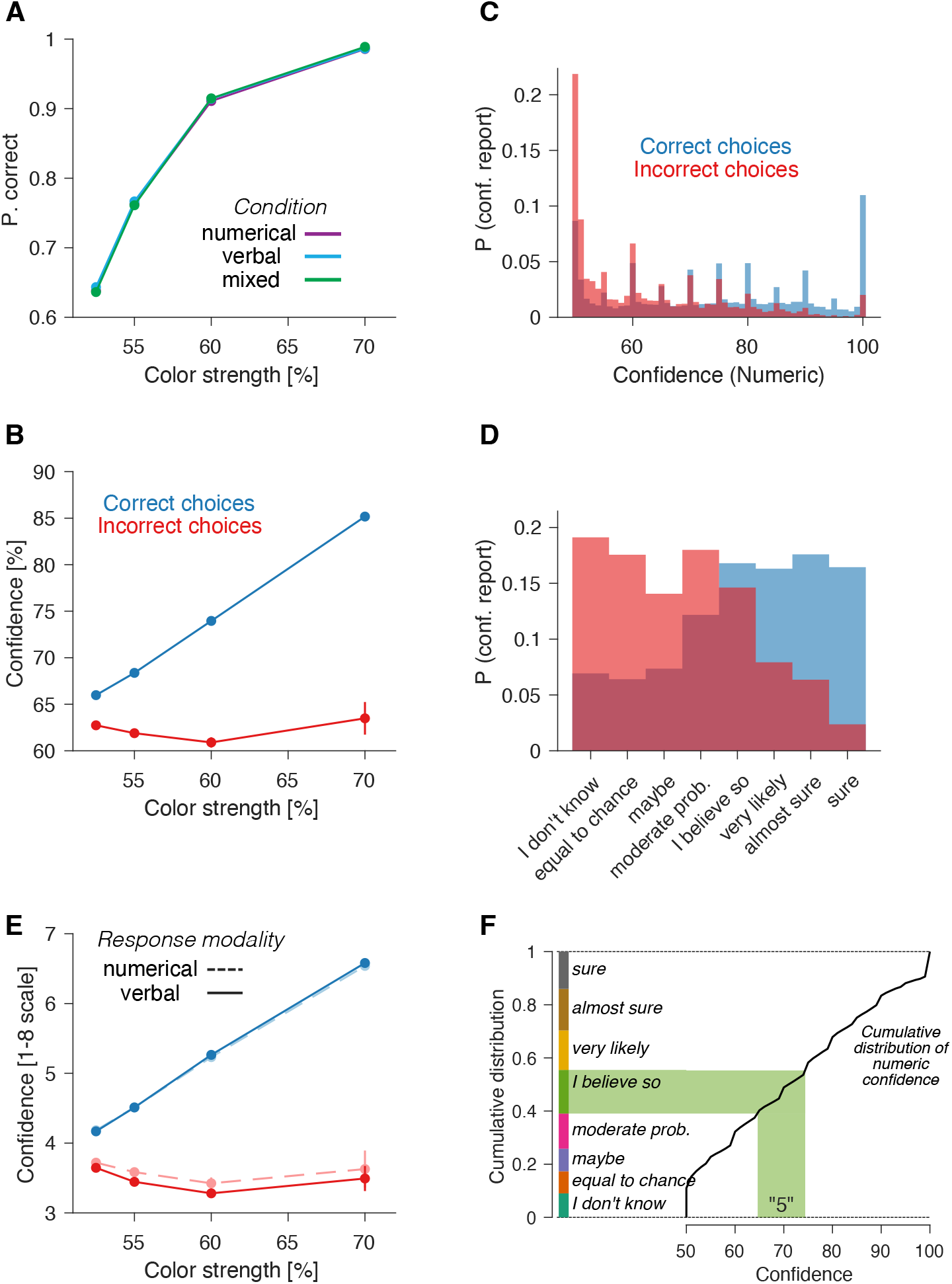
Initial Choice and Confidence Reports. **(A)** Proportion of correct choices as a function of color strength, calculated separately for the Numerical Only, Verbal Only, and Mixed conditions. **(B)** Mean confidence (Numerical Only condition) as a function of color strength, computed separately for correct and incorrect choices. Error bars (mostly smaller than the data points) represent standard errors. **(C)** Distribution of numeric confidence reports (Numerical Only condition), grouped by whether the choice was correct (blue) or incorrect (red). Confidence reports could range from 50 to 100 in integer values. **(E)** Distribution of verbal confidence reports conditioned on choice accuracy. The confidence reports were obtained from the Verbal Only and Mixed conditions. **(F)** Mean confidence as a function of color strength, shown separately for correct (blue) and incorrect (red) decisions, and for verbal (solid curves) and numeric (dashed curves) reports. Verbal confidence reports were mapped to a 1–8 scale, rank-ordered by the mean probability assigned in the mapping task. Numeric confidence reports were mapped to the same scale using the transformation described in panel F. **(G)** Mapping between numerical and verbal confidence reports. Numerical confidence values were matched to verbal categories such that the fraction of trials within each category were equal across the two reporting modalities.

Numerical confidence reports followed expected trends: higher confidence for correct than incorrect choices, and increased confidence with stronger color strength for correct choices (Fig. 3B). For incorrect choices, confidence showed less dependence on color strength (*β*_1_ = 110 ± 1.5, *β*_2_ = −10 ± 7, Eq. 4). Participants often reported confidence in multiples of five, which may be indicative of coarse granularity in the internal sense of confidence (Fig. 3C) (Ott, 2021; Lisi et al., 2021; Ais et al., 2016).

Verbal confidence reports were also informative about choice accuracy. Participants tended to use high-confidence expressions (e.g., “Sure”) for correct decisions and lower-confidence expressions (e.g., “Moderate probability”) for incorrect ones (Fig. 3D). We ranked the verbal expressions by the mean probabilities assigned to them in the mapping experiment (Fig. 2). We averaged the ranks (from 1 to 8) of the verbal expressions as a function of color strength, separately for correct and incorrect decisions. Similar to the numerical reports, verbal confidence increased with color strength for correct choices and was less dependent on color strength for errors (Fig. 3E, solid traces).

To compare verbal and numerical confidence reports on a common scale, we grouped the numerical confidence reports into eight categories, each one corresponding to a different verbal expression. Higher numerical values were assigned to higher-ranked categories, and the proportion of trials in each category matched the frequency with which the corresponding verbal expression was used (Fig. 3F). For instance, if the expression “Sure” was used in *y*% of the trials, the top *y*% of numerical confidence values were assigned to the category representing that expression.

This transformation made evident the consistency between verbal and numerical confidence reports. For correct decisions, the average confidence was nearly identical across response modalities (Fig. 3E) (*p* = 0.28; ℋ_0_ : *β*_2_ = 0; Eq. 5). However, for incorrect decisions, numerical reports were slightly but significantly higher than verbal reports (*p* = 0.005; ℋ_0_ : *β*_2_ = 0; Eq. 5). As a result, the difference in confidence between correct and incorrect choices is larger for verbal expressions, indicating potentially greater metacognitive sensitivity. In the following section, we evaluate this assertion quantitatively, using a Receiver Operating Characteristic (ROC) analysis.

### The Metacognitive Sensitivity of Verbal and Numerical Confidence Reports

We quantified how well initial confidence reports distinguish between correct and incorrect choices using the area under the ROC curve (AUC_conf_; Fig. 4A). The AUC_conf_ represents the probability that, given two confidence values—one randomly drawn from a trial with a correct choice and the other from a trial with an incorrect choice—the higher confidence value is associated with the correct trial. This metric is derived from the distributions of confidence reports for correct and incorrect choices (Fig. 3B,D).

**Figure 4.**
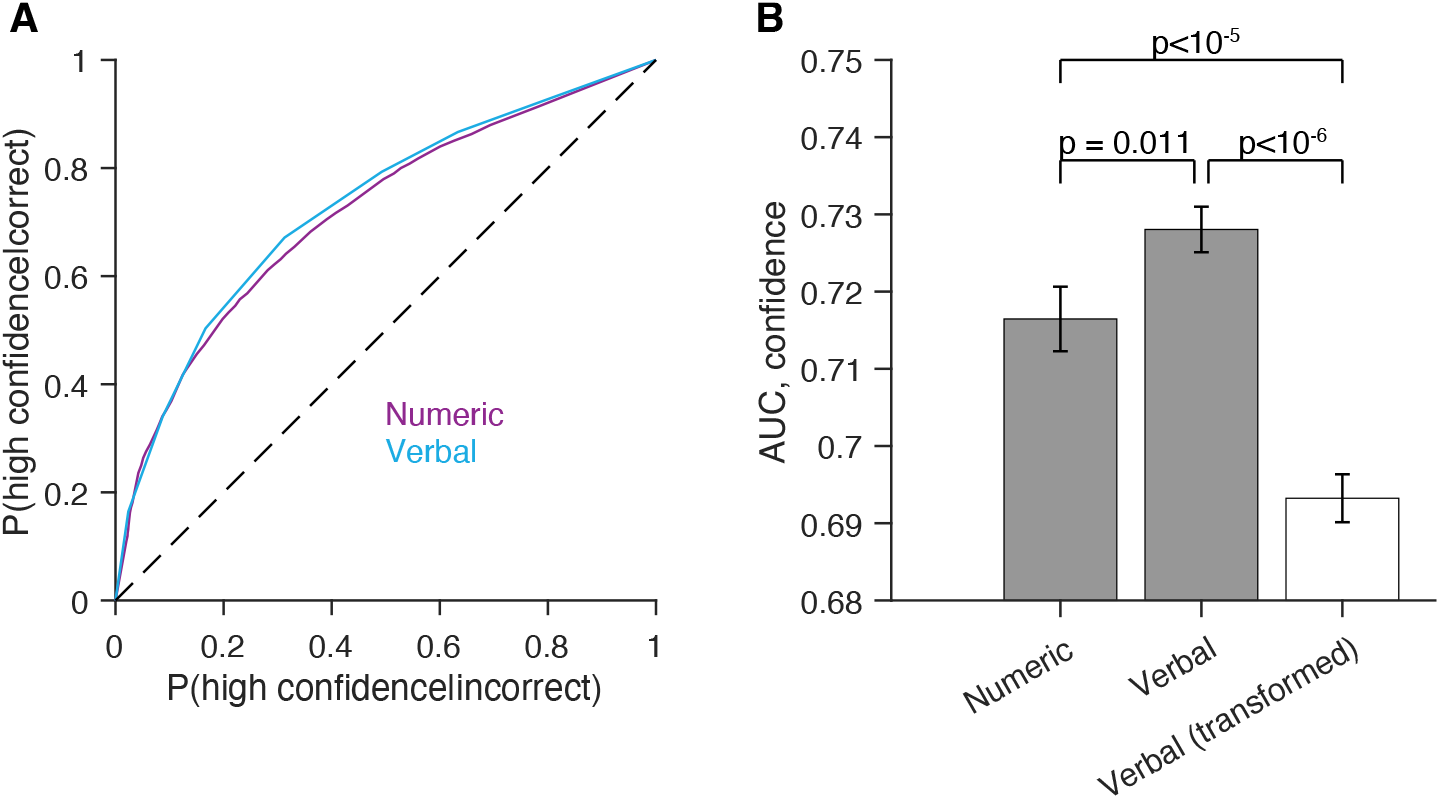
Metacognitive Sensitivity of Numerical and Verbal Confidence Reports. **(A)** ROC curves derived from the distribution of verbal and numerical confidence reports (Fig. 3C,D). Verbal confidence reports were obtained from the Verbal Only and Mixed conditions. **(B)** Area under the ROC curve (AUC) for verbal and numerical confidence reports. AUC_conf_ is significantly higher for verbal reports. The rightmost bar represents the AUC obtained when verbal reports are converted into probability values based on explicit probability estimates from the mapping experiment (Fig. 2). Error bars indicate standard error (s.e.). P-values and s.e. were computed using a bootstrap analysis.

We compared AUC_conf_ for initial confidence reports in verbal and numerical modalities and found that verbal reports yielded a significantly higher AUC_conf_ than numerical reports (Fig. 4B; p = 0.011, bootstrap). This result suggests that verbal confidence reports provide a slightly but significantly better distinction between correct and incorrect choices. To validate this finding, we conducted a logistic regression analysis (Rausch and Zehetleitner, 2017). Choice accuracy (correct vs. incorrect) is predicted from variables including color strength, confidence, and response modality (verbal or numerical) and interaction terms (Eq. 1). The key covariate is the interaction term between confidence and response modality: a significant effect of choice accuracy would indicate a difference in confidence resolution between verbal and numerical reports. Consistent with the AUC_conf_ analysis, the regression showed that confidence resolution was significantly higher for verbal reports (*β*_4_ = −0.026 ± 0.01; p = 0.02; ℋ_0_: *β*_4_ = 0; Eq. 1).

Since most decision models rely on numerical probabilities (e.g., Kahneman, 1979; Savage, 1954), it is plausible that verbal confidence reports are interpreted as numerical probability values. If so, the probability value assigned to each verbal expression in the mapping task (Fig. 2) should be more informative about choice accuracy than the verbal expression itself. To test this, we transformed verbal confidence reports using the explicit probability estimates provided by each participant in the mapping experiment (Fig. 2) and recalculated AUC_conf_ based on the resulting probability values. Contrary to the hypothesis, confidence resolution was substantially lower than that obtained from the actual verbal reports (Fig. 4B, open bar). This suggests that verbal reports contains more information about decision accuracy than their numerical interpretation derived from explicit probability estimates.

### Decision Revisions are Informed by Advice Regardless of Report Modality

The finding that verbal confidence reports exhibit greater metacognitive sensitivity than numerical ones challenges the belief that verbal expressions are inherently more ambiguous or imprecise. However, this interpretation hinges on the experimenters’ evaluation of verbal reports’ ability to discriminate between correct and incorrect choices. We assessed whether participants themselves could interpret others’ verbal confidence reports and use them to improve their own decisions. To this end, we analyzed the likelihood that participants change their mind about their choice after seeing the advice, as a function of the initial and advice confidence.

We analyzed trials where participants’ initial choice differed from the advice choice, as these are the trials where changes of mind might occur. For these trials, we calculated the likelihood of a change of mind as a function of initial confidence and the confidence conveyed in the advice, separately for verbal and numerical reporting modalities. Changes of mind were more likely when (*i*) participants’ initial confidence was low (*β*_2_ = −0.37 ± 0.019, Eq. 6) and (*ii*) the advice confidence was high (*β*_3_ = 0.25 ± 0.017, Eq. 6)(Fig. 5). These results show that both initial confidence and advice confidence influence the likelihood of revising an initial choice.

**Figure 5.**
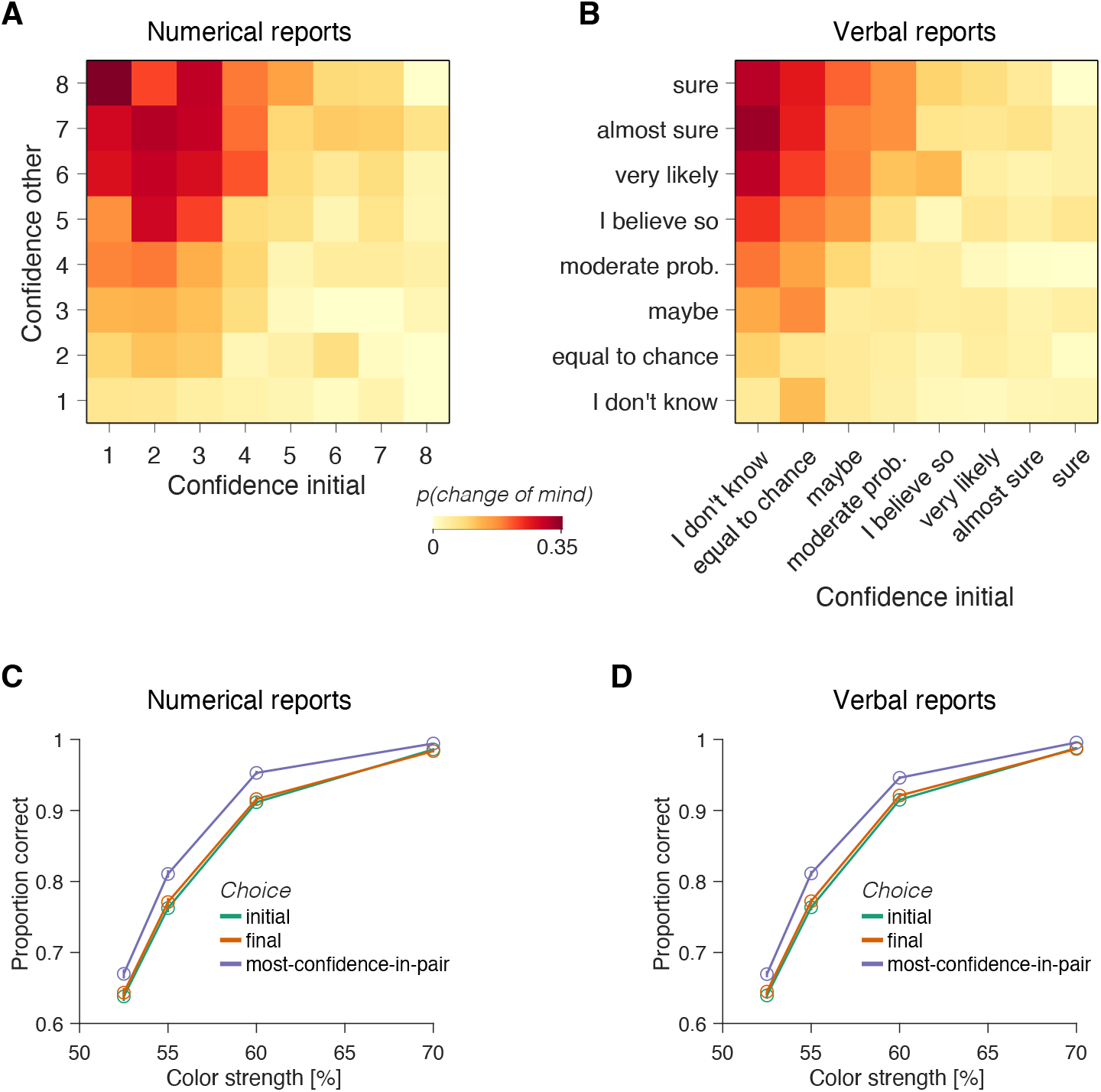
Impact of Advice on Decision Revisions Across Response Modalities. **(A)** Proportion of trials with a change of mind as a function of initial confidence (x-axis) and advice confidence (y-axis). The analysis includes only trials where the participant’s initial choice and the advice choice differed (i.e., trials where changes of mind were possible). Numerical confidence reports were categorized as detailed in Fig. 3F. **(B)** Same as panel A, but for verbal confidence reports (conditions Verbal Only and Mixed). **(C)** Proportion of correct choices as a function of color strength. The green curve represents accuracy based on initial choices, while the orange curve reflects accuracy based on final choices. The purple curve illustrates hypothetical accuracy if final choices were determined by always selecting the option with the highest confidence, whether from the initial report or the advice. Only trials with numerical confidence reports (Numerical Only condition) were included. **(D)** Same as panel C, but using trials with verbal confidence reports (Verbal Only and Mixed conditions).

The regression analysis revealed no significant effect of the report modality on the likelihood of a change of mind about the choice. Neither the interaction between self-confidence and reporting modality (*p* = 0.13, ℋ_0_: *β*_5_ = 0, Eq. 6) nor the interaction between advice confidence and reporting modality (*p* = 0.22, ℋ_0_: *β*_6_ = 0,Eq. 6) significantly impacted the probability of a change of mind. These findings indicate that numerical and verbal confidence reports were interpreted similarly by participants when considering whether to revise their initial choice.

### Suboptimal Use of Advice to Revise Initial Choices

Although advice influenced participants’ final decisions, they changed their minds less often than would have been optimal for this task. Given that both initial self-reports and advice provide comparable information about decision accuracy, they should, in principle, be weighted equally in final decisions. However, this was not the case: even when initial confidence was at its lowest (“I don’t know”) and advice confidence was at its highest (“Sure”), participants changed their minds in fewer than 40% of cases (Fig. 5A-B).

The suboptimal use of the advice is further highlighted by a simple strategy that combines the two choices and outperforms participants’ accuracy. If participants consistently selected the option associated with the highest confidence—whether from their own initial report or the advice—their accuracy would have been significantly higher than observed (Fig. 5C-D; *β*_2_ = 0.47 ± 0.05; *p* < 10^−8^; ℋ_0_ : *β*_2_ = 0; Eq. 7). The tendency to underweight the opinions of others aligns with findings from previous studies (e.g., Yaniv, 2004).

### Stability of Metacognitive Sensitivity Over Trials

Our experiment involved a large number of participants completing only eight trials each, enabling an analysis of how confidence and metacognitive sensitivity evolves over trials. Accuracy increased significantly across trials (Fig. 6A), as did confidence, the difference in confidence between correct and incorrect decisions, and the AUC_conf_ (Fig. 6A-C).

**Figure 6.**
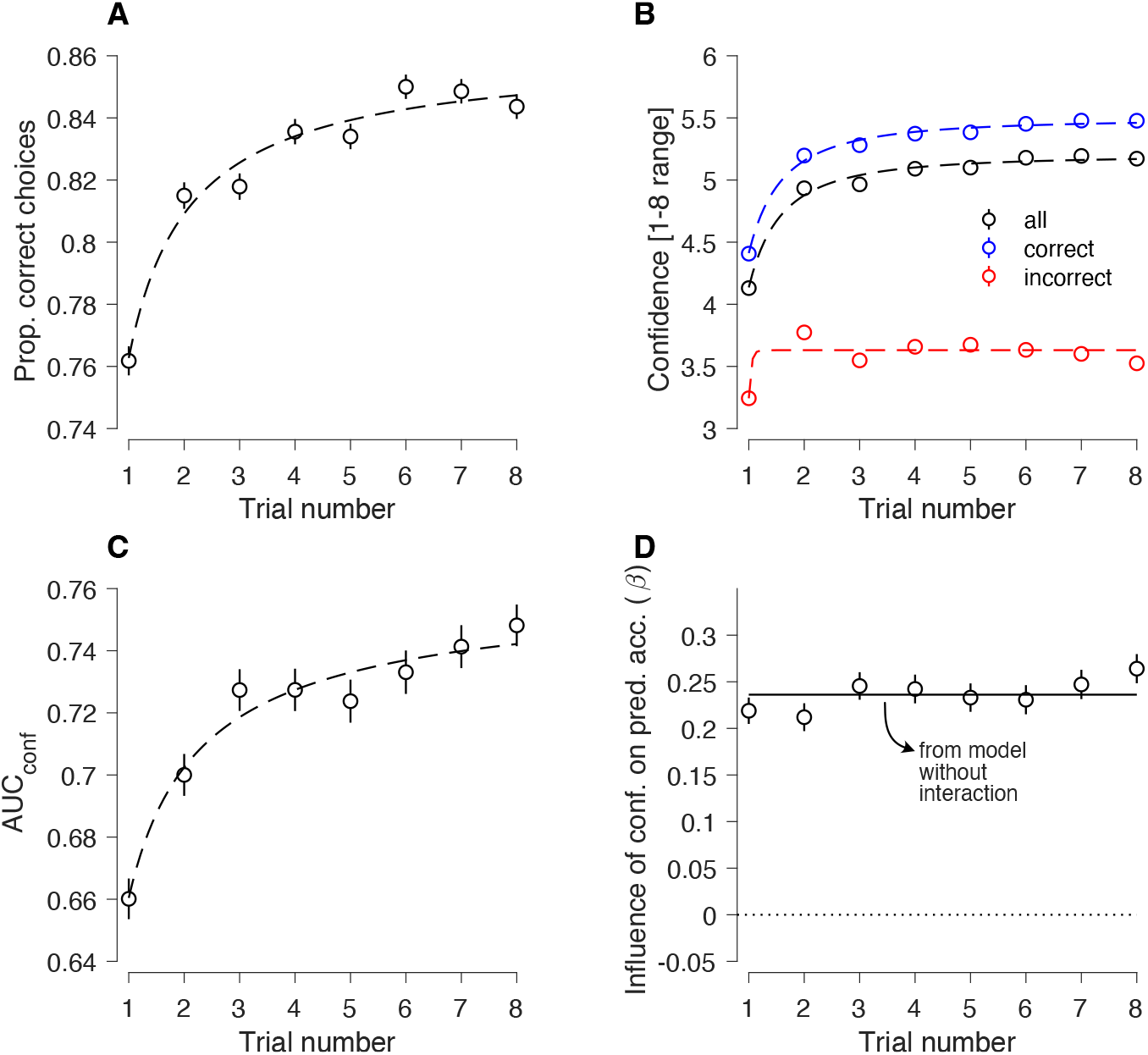
Consistency of Metacognitive Sensitivity Over Successive Trials. **(A)** Proportion of correct choices as a function of trial number. Error bars represent s.e., and dashed lines are fits of a modified power-law function with an offset, *F* (*x*) = *a* − *b* (*e*^−*cx*^*x*^−*d*^), where {*a, b, c, d*} are fit parameters. The same convention applies to the other graphs in this figure. **(B)** Mean reported confidence as a function of trial number for correct (blue), incorrect (red), and all (black) trials. Numerical confidence reports were binned as described in Fig. 3F to allow combination with verbal confidence reports. **(C)** AUC_conf_ as a function of trial number. **(D)** Capacity of confidence reports for distinguishing between correct and incorrect choices, estimated using logistic regression. Data points represent estimates from a model with a separate regression coefficient for confidence for each trial number, while the horizontal line represents estimates from a model without an interaction term between confidence and trial number.

Because choice accuracy varied across trials, we assessed metacognitive sensitivity using logistic regression instead of AUC_conf_. The regression model was designed to predict whether a choice was correct or incorrect and included interaction terms for color strength, confidence, and trial number (Eq. 2). Sensitivity to color strength increased significantly over trials, consistent with the observed improvement in accuracy (*p* < 10^−8^, likelihood-ratio test, ℋ_0_: *β*_2⋅_ = 0, Eq. 2). However, there was no evidence that the ability of confidence to predict accuracy changed over trials. A comparison of nested models with and without the interaction between confidence and trial number favored the model without the interaction (Fig. 6D; *p* = 0.29, likelihood-ratio test, ℋ_0_ : *β*_4⋅_ = 0, Eq. 2).

Although these analyses aggregate confidence reports across conditions, similar results were observed when verbal and numerical confidence reports were analyzed separately (Figs. S1 and S2). These findings indicate that confidence remains equally informative about decision accuracy in both early and late trials, despite notable changes in overall accuracy and confidence.

### Grounded Translation of Verbal Expressions of Uncertainty

The experiments were conducted in Spanish, and the verbal probability expressions were translated literally into English for the reader. However, because we are studying verbal expressions that have a numerical counterpart (Fig. 2), an alternative translation approach is possible. Instead of relying solely on literal translation, we can identify English expressions whose meaning, in terms of their mapping onto a numerical scale, most closely aligns with the Spanish expressions. We refer to this approach as grounded translation.

Mauboussin and colleagues conducted an experiment similar to our mapping task (Fig. 2) in English (Mauboussin and Mauboussin, 2018). For each verbal expression in our study, we identified the closest equivalent in Mauboussin’s experiment. To determine similarity, we measured the Kullback-Leibler (KL) divergence between the distributions of numerical reports associated with each verbal expression.

Fig. 7 presents the closest English equivalent for each Spanish expression according to this metric. For comparison, we also include the literal translation obtained by prompting a large language model (ChatGPT-4o) to select the most appropriate English expression for each Spanish term from the available options. The high consistency between literal and grounded translations validates the use of literal translations, while illustrating the viability of a more nuanced, data-driven translation approach.

**Figure 7.**
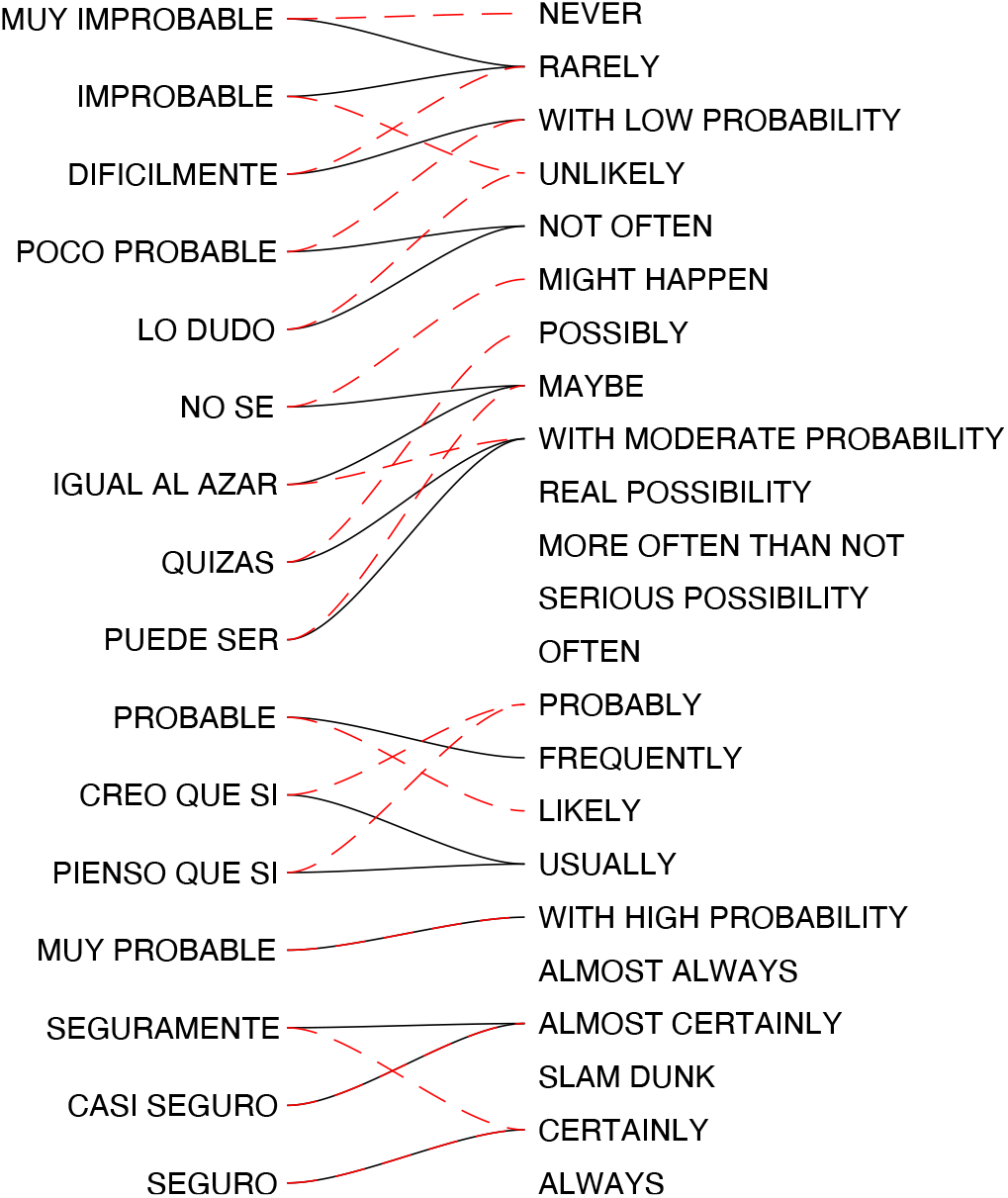
Grounded translation of verbal expressions. For each Spanish verbal expression used in the mapping task (Fig. 2), we identified the most similar expression from the list in Mauboussin and Mauboussin (2018)’s experiment. The black lines represent the matches obtained by minimizing the Kullback-Leibler (KL) divergence between the distributions of numerical reports from both experiments. The dashed red lines indicate the matches generated by prompting a large language model (ChatGPT-4o). Spanish and English expressions are ordered from top to bottom by the average probability assigned to each.

## Discussion

Effectively communicating confidence is crucial across various domains, including science, policy-making, and interpersonal collaboration. Confidence reflects a subjective belief in the accuracy of a decision and influences both decision-making and the integration of others’ opinions. Our study examined the metacognitive sensitivity of verbal and numerical confidence reports—their ability to distinguish between correct and incorrect decisions. We assessed confidence both directly, through explicit reports, and indirectly, by measuring the weight decision-makers assigned to others’ judgments when revising an initial choice.

Verbal confidence reports—treated as an ordinal variable—exhibited slightly higher metacognitive sensitivity than numerical reports. That is, verbal reports provided a better indication of decision accuracy than numerical ones. Using a simple binary perceptual decision task, we parametrically controlled decision difficulty, revealing that both verbal and numerical confidence reports followed a similar pattern as a function of decision difficulty and accuracy. Following an error, verbal confidence reports were lower than numerical ones, which explains the higher metacognitive sensitivity of verbal than numerical reports. Further, participants effectively integrated their own confidence with another participant’s advice, regardless of whether confidence and advice were conveyed verbally or numerically. These results challenge the common assumption that verbal expressions are inherently more ambiguous or less precise than numerical ones when communicating degrees of uncertainty (Budescu et al., 2014; Brun and Teigen, 1988; Lichtenstein and Newman, 1967; Ott, 2021; Mauboussin and Mauboussin, 2018; Beyth-Marom, 1982; Mosteller and Youtz, 1990).

The large number of participants in our experiment allowed us to evaluate metacognitive sensitivity on a trial-by-trial basis. Our results show that while choice accuracy and confidence increased significantly with task experience, metacognitive sensitivity remains stable (Fig. 6). This pattern held regardless of whether confidence was reported verbally or numerically. In other words, while participants improved at making perceptual decisions with experience, their ability to assess their own choice accuracy did not, suggesting that different learning processes underlie these two types of judgments. The stability of metacognitive sensitivity across trials supports prior studies that assessed metacognitive sensitivity by aggregating data from multiple trials of individual participants (e.g., Ais et al., 2016). Note that, although AUC_conf_ increased substantially over trials, this metric is unsuitable for comparing conditions with different levels of choice accuracy, leading us to rely instead on a logistic regression-based measure (Maniscalco and Lau, 2012; Rausch and Zehetleitner, 2017).

The high metacognitive sensitivity of verbal confidence reports aligns with the idea that verbal and numerical expressions capture different forms of uncertainty (Ülkümen et al., 2016). A useful distinction, introduced by Fleming (2024), differentiates between *world-centered uncertainty* —such as predicting whether it will rain tomorrow—and *self-centered uncertainty* —such as confidence in one’s own decisions or actions. Most studies comparing verbal and numerical reports focus on world-centered uncertainty, such as predicting the outcome of a sporting event (Erev and Cohen, 1990), estimating the probability of a geopolitical event (Beyth-Marom, 1982), or assessing the likelihood that a dart will land in a shaded area on a spinner (Budescu et al., 1988). In contrast, our study examines confidence reports related to the accuracy of a perceptual decision—judgments based on internal, unobservable factors that cannot be directly interpreted in terms of frequentist probabilities. While speculative, it is possible that verbal expressions are particularly well-suited for conveying self-centered confidence judgments, whereas numerical scales may be more effective for communicating world-centered uncertainties.

## Methods

### Experimental Design

The experiment consisted of four stages, only a subset of which are analyzed in this study:

- **Stage 1: Mapping task** (Fig. 2). Participants reported the number (0–100 range) that best represents the probability value of each of 16 verbal expressions of uncertainty.
- **Stage 2: Mapping verbal expressions to probability ranges** (Fig. S4A). Participants assigned a probability range to each of the 16 verbal expressions used in Stage 1 by adjusting two sliders: one for the lower bound and another for the upper bound.
- **Stage 3: Conjunction of expressions of uncertainty** (Fig. S4B). Participants were shown pairs of uncertainty expressions, which could be numerical and/or verbal (e.g., *Maybe* and 60%, *Sure* and *Likely*, or 70% and 60%). They then selected the numerical probability that best represented the likelihood of an event described by both expressions.
- **Stage 4: Color Experiment** (Fig. 1).

Our study focuses only on the data from Stages 1 and 4.

### Participants

The experiment included 8,308 participants (mean age: 29 years). 40.42% of participants completed the experiment on a desktop computer, while the rest used a mobile device; data from both were combined for analysis as no notable difference was observed. No participants who completed the experiment were excluded from the analyses.

### Main Task

In the main task, participants judged whether a circular aperture contained more red or blue dots (Fig. 1A). The *color strength* was defined as the proportion of dots belonging to the majority color and took one of four possible values: 52.5%, 55%, 60%, or 70%. Each participant completed two trials at each color strength—one where blue was the majority and one where red was the majority—resulting in a total of eight trials per participant.

Numerical confidence reports were made using a horizontal scale with 50% labeled at the midpoint and 100% at the rightmost end. The portion of the scale below 50% was visible but not clickable and was displayed in light gray (Fig. 1B). When participants clicked on the scale, a circular marker appeared, displaying the closest integer between 50 and 100 corresponding to the selected position. They could revise their response and adjust their confidence as many times as desired before pressing the “Next” button to proceed.

Verbal confidence reports were collected by selecting an expression from a predefined list of eight options. These expressions were ordered from lowest to highest confidence based on the average numerical values assigned to them in the mapping experiment (Fig. 2), with the final ordering shown in Fig. 3F.

### Color Stimulus

The color stimulus consisted of *N*=40 dots, randomly and uniformly distributed within a circular aperture. To prevent overlap, any dots that intersected with others were resampled as many times as necessary.

The aperture’s diameter was set to the smallest of three values: 300 pixels, 0.8*w*, or 0.4*h*, where *w* and *h* represent the screen’s width and height, respectively. For nearly all participants the aperture was 300px in diameter. Each dot’s diameter was 2% of the aperture’s diameter (i.e., 6px for most participants).

### Mapping Confidence Expressions to Probability Estimates

In the *mapping* task, completed before the main task, participants were presented with 16 verbal expressions of uncertainty in random order and asked to indicate the probability value that best represented each expression using a numerical scale (Fig. 2). The scale ranged from 0 to 100, with these values explicitly marked at the endpoints. When participants clicked on the scale, a circular marker appeared at the selected position, displaying the corresponding probability. They could adjust the marker’s position as many times as desired, with the final selection recorded as their response.

### Data Analysis

#### Rank-ordering of the verbal expressions of probability

We ranked the verbal expressions used for confidence reporting based on the average probability values assigned to them in the mapping task (Fig. 2). An alternative way to rank verbal expressions, that does not rely on the mapping task, is to sort them post hoc based on the proportion of correct responses observed when each expression was used. However, this approach risks artificially inflating the metacognitive resolution of verbal expressions, as it does not guarantee that the ranking reflects how participants inherently interpret these terms. In our data, however, both methods produce a similar ordering (Fig. S3).

#### Discretization of Numerical Confidence Reports

For some analyses, we discretized the numerical confidence reports into eight categories, each corresponding to one of the verbal expressions used to report initial confidence in the experiment. To do this, we calculated the proportion of trials in which each verbal expression was chosen as the initial confidence (Verbal Only and Mixed conditions). We then grouped the initial numerical confidence reports from the Numerical Only condition into eight categories so that the proportion of trials in each category matched that obtained from the verbal confidence reports (Fig. 3F). This mapping establishes a set of criteria on the numerical confidence scale that separates the different confidence categories.

#### Metacognitive Sensitivity

To quantify metacognitive sensitivity, a type-2 ROC analysis was performed using confidence reports for correct and incorrect choices (Clarke et al., 1959). The area under the type-2 ROC curve (AUC_conf_) was used as a measure of metacognitive sensitivity. To assess whether one AUC_conf_ value was significantly greater than another (e.g., comparing verbal and numerical reports, Fig. 4), a bootstrap approach with *N* = 5000 samples per condition was used. Significance was determined by computing the proportion of bootstrap samples where the tested condition was satisfied. The bootstrap procedure preserved the proportion of trials across difficulty levels.

The ROC analysis (Fig. 4) was validated using a logistic regression model:

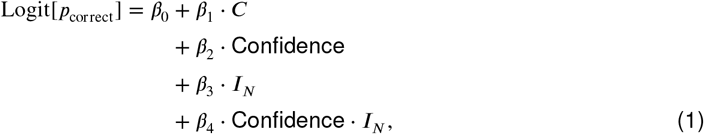

where *C* denotes color strength, and *I*_*N*_ is an indicator variable set to 1 for the Numerical Only condition and 0 otherwise. Confidence values range from 1 to 8; for verbal reports, these correspond to the rank ordering of the verbal expressions based on their assigned probability in the mapping task, while for numerical reports, confidence values were binned into eight discrete categories as detailed in Fig. 3F.

A logistic regression model was used to examine whether the ability of confidence to predict choice accuracy varied across trials:

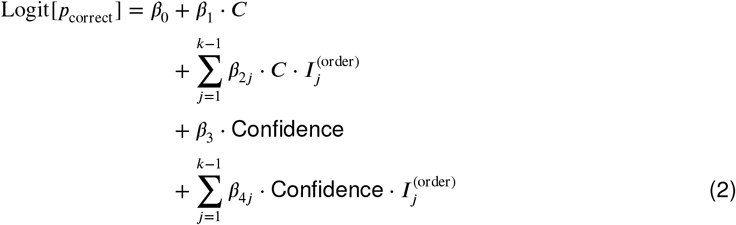

where 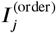 is an indicator variable set to 1 for trial *j* and 0 otherwise, and *k* = 8 represents the total number of trials per participant. The data points in Fig. 6D correspond to the regression coefficient quantifying the influence of confidence on the accuracy prediction, given by (*β*_3_ + *β*_4*j*_) for trials 1 to 7 and *β*_3_ for trial 8. The solid line in the same graph represents the model fit when the *β*_4⋅_ terms are omitted. A likelihood ratio test was used to compare the model with and without the *β*_4⋅_ terms. The comparison favored the simpler model. Numerical confidence values were discretized into eight categories, as described above, to be able to combine them with the verbal reports on a common scale. However, in Fig. S2, this discretization was not applied, as the analysis includes only trials with numerical confidence reports.

### Statistical Analysis

Logistic regression was used to assess the influence of color strength and response modality on choice accuracy:

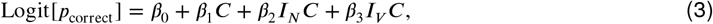

where *C* represents color strength, and *I*_*N*_ and *I*_*V*_ are indicator variables for the Numerical Only and Verbal Only conditions respectively. To evaluate the effect of response modality on accuracy, a likelihoodratio test was performed to compare nested models with and without the *β*_2_ and *β*_3_ terms.

Linear regression was used to examine how confidence depends on color strength and choice accuracy:

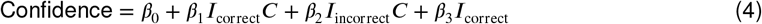

where *I*_correct_ and *I*_incorrect_ are indicator variables for correct and incorrect choices, respectively. The significance of the regression coefficients was assessed using a t-test based on the standard errors of the estimated coefficients.

We also used linear regression to determine if confidence depends on response modality (Fig. 3E):

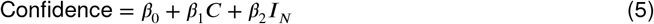

The model was fit independently for correct and incorrect choices.

To examine the factors that influence the likelihood of decision-makers changing their choice between the initial and final report, the following logistic regression model was applied:

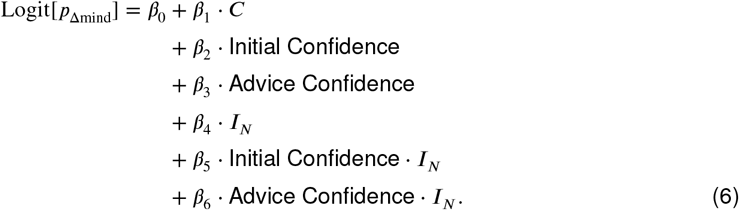

The effects of initial and advice confidence were assessed through the coefficients *β*_2_ and *β*_3_, respectively, while their interaction with response modality was captured by the *β*_5_ and *β*_6_ terms. “Initial Confidence” and “Advice Confidence” range from 1 to 8. Numerical confidence reports were discretized into these categories, as shown in Fig. 3F, to incorporate into the same regression model as verbal expressions.

To evaluate whether choice accuracy could have been improved by selecting the decision made with the highest confidence—either the participant’s initial choice or the advice received—we applied the following logistic regression model:

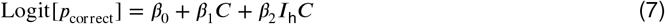

The dependent variable combines the accuracy of the participant’s final decision with the hypothetical accuracy of selecting the most confident choice. The indicator variable *I*_h_ was set to 1 for hypothetical choices and 0 for the participant’s actual choices.

#### Spanish-English Translation

The *grounded* translations were determined by identifying, for each Spanish verbal expression, the corresponding English expression from Mauboussin and Mauboussin (2018) that minimized the Kullback-Leibler (KL) divergence:

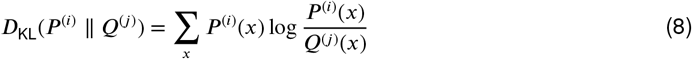

where *P*^(*i*)^(*x*) represents the proportion of trials in which participants assigned a value of *x* to the Spanish expression *i* in our experiment, and *Q*^(*j*)^(*x*) is the corresponding proportion for the English expression *j* in Mauboussin and Mauboussin (2018). The numerical scale was discretized into 100 bins. To avoid division by zero in Eq. 8, each bin’s response count was incremented by 1 before calculating the proportion of responses in each bin (Laplace smoothing).

To obtain the *literal* translations, we prompted chatGPT-4o (February 2025) with the following text:

For each of these verbal expressions of uncertainty in spanish, {‘Seguro’, ‘Casi seguro’, ‘Seguramente’,’ Muy probable’, ‘Pienso que si’, ‘Creo que si’, ‘Probable’, ‘Puede ser’, ‘Quizas’, ‘Igual al azar’, ‘No se’, ‘Lo dudo’, ‘Poco probable’, ‘Dificlmente’,’Improbable’, ‘Muy improbable’}, find the verbal expression in english that better reflects its meaning, out of the ones in this list: {‘Always’, ‘Certainly’, ‘Slam dunk’, ‘Almost certainly’, ‘Almost Always’, ‘with high probability’, ‘usually’, ‘likely’, ‘frequently’, ‘probably’, ‘often’, ‘serious possibility’, ‘more often than not’, ‘real possibility’, ‘with moderate probability’, ‘maybe’, ‘possibly’, ‘might happen’, ‘not often’, ‘unlikely’, ‘with low probability’, ‘rarely’, ‘never’}.

## Ethics statement

This study is based on anonymized data collected through an online experiment conducted via *El Gato y la Caja*, a Spanish-language science communication platform based in Argentina (https://elgatoylacaja.com). The experiment was programmed and hosted by the platform. Participation was voluntary and unpaid, and participants provided informed consent prior to participation. No personally identifiable information was included in the dataset shared with the authors.

A previous analysis of the data was published at https://elgatoylacaja.com/quizas-quizas-quizas.

## Acknowledgments

We are deeply grateful to the *El Gato y La Caja* team for their essential contributions to the development and communication of this experiment and for sharing the data with us. Special thanks to Vicky Milano, Juan Cuiule, Laura González, Rocco Di Tella, Pablo González, Valeria Sanabria, Emma Coso, Azul Damadián, Ezequiel Calvo, Belén Kakefuku, Juan Manuel Garrido, Javier Goldschmidt, and Florencia Gonzalez. We also extend our gratitude to *Comunidad Gato*, whose participation and support helped broaden the study’s reach. We also thank Pablo Barttfeld and Guillermo Solovey for helpful discussions.

## Author contributions

F.A.H. and A.Z. conceived and designed the research.

F.A.H. and A.Z. conducted the research.

A.Z. analyzed the data.

A.Z. wrote the original draft.

All authors revised and edited the manuscript.

## Supplemental information

**Figure S1.**
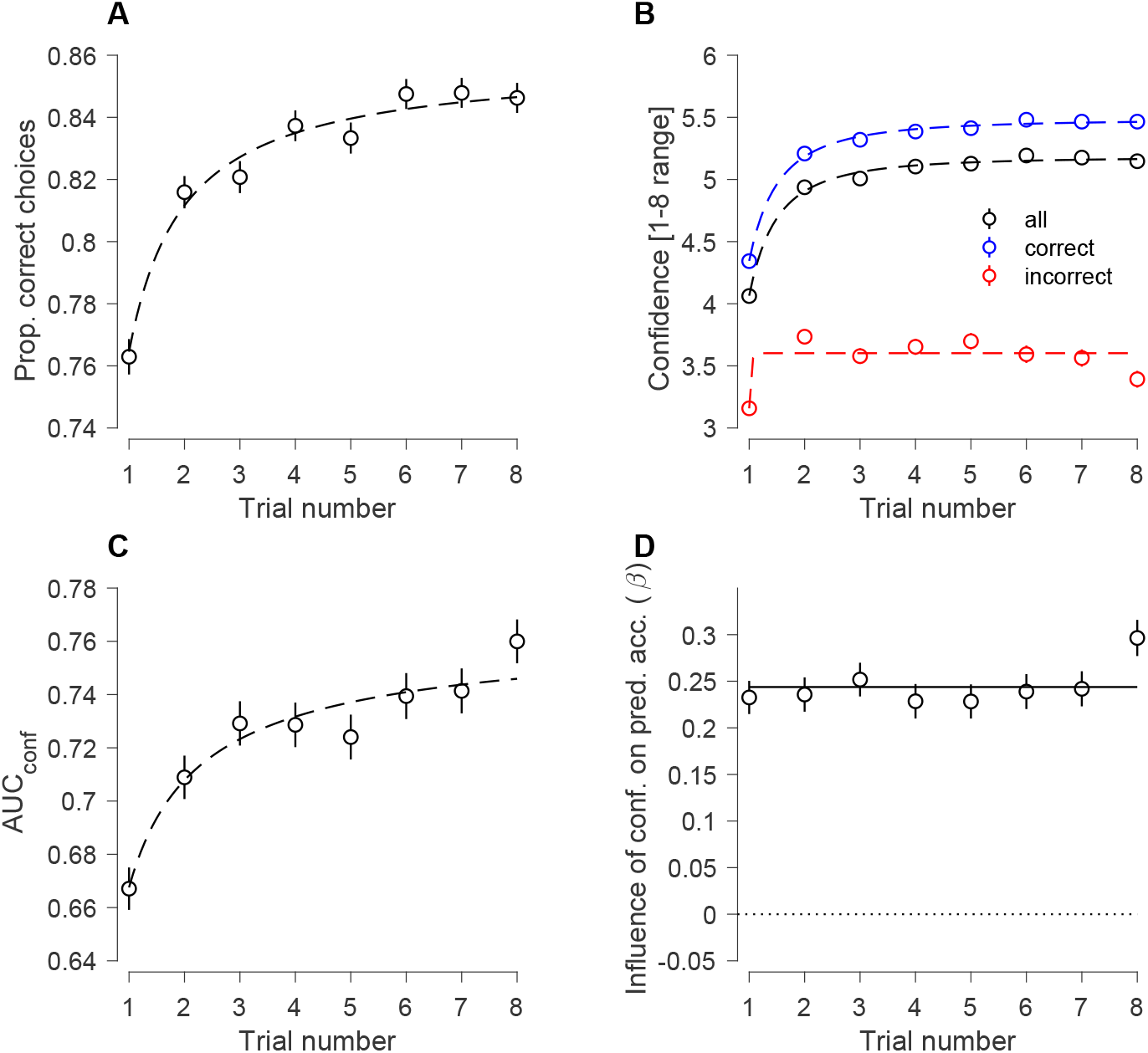
Changes in Verbal Confidence Reports Across Trials. Same as Fig. 6, using only trials with verbal confidence reports (Verbal Only and Mixed conditions).

**Figure S2.**
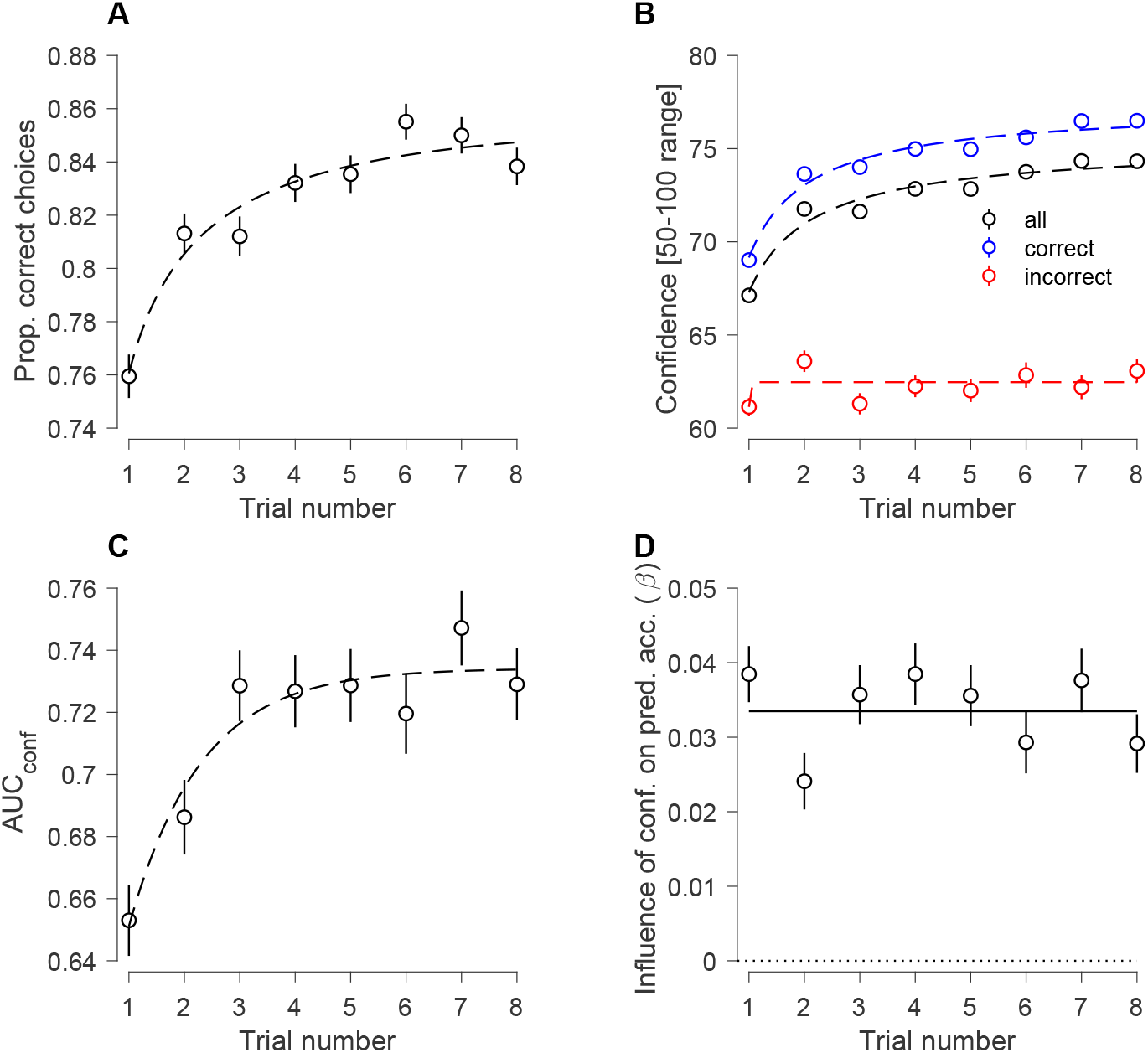
Changes in Numerical Confidence Reports Across Trials. Same as Fig. 6, using only trials with numeric confidence reports (Numerical Only condition).

**Figure S3.**
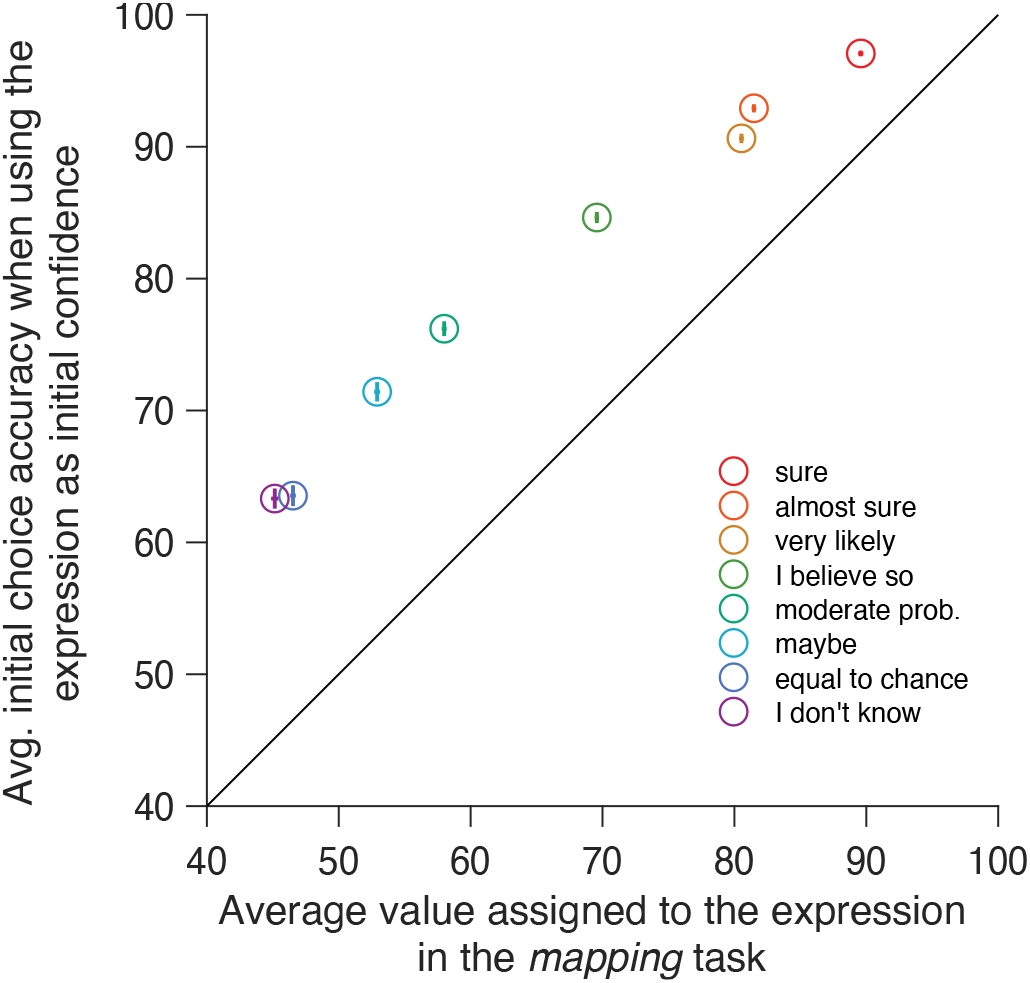
Relation Between Assigned Probability Values and Choice Accuracy. Average probability value assigned to each verbal expression in the mapping task (x-axis) plotted against the proportion of correct initial color choices when participants selected that expression to report their confidence. Error bars indicate s.e.

**Figure S4.**
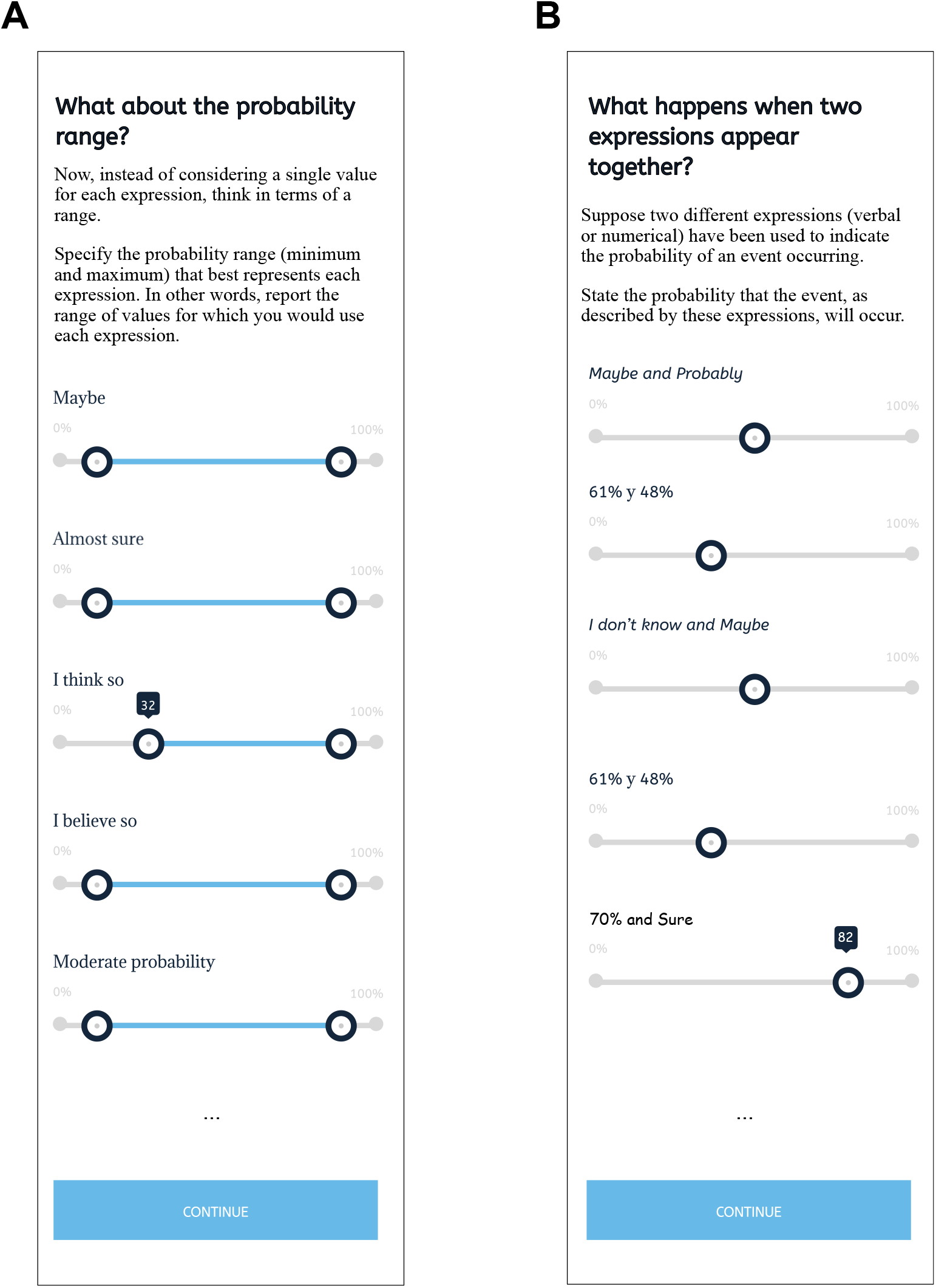
Assessing Probability Ranges and Combining Uncertainty Expressions. Participants completed two additional tasks during the same session, following the mapping task (Fig. 2) and preceding the color discrimination task (Fig. 1). Data analyses from these tasks will be reported elsewhere. The figure panels closely resemble what participants observed. **(A)** Range-estimation task: Participants specified the range of values (minimum and maximum) that best captured the meaning of each expression. **(B)** Expression-conjunction task: Participants were presented with pairs of uncertainty expressions (numerical and/or verbal) and were asked to report the probability of occurrence for the event described by these expressions.

